# Phase *i* trials in melanoma: A framework to translate preclinical findings to the clinic

**DOI:** 10.1101/015925

**Authors:** Eunjung Kim, Vito W. Rebecca, Keiran S.M. Smalley, Alexander R.A. Anderson

## Abstract

**Abstract:** We present a, mathematical model driven, framework to implement virtual or imaginary clinical trials (phase *i* trials) that can be used to bridge the gap between preclinical studies and the clinic. The trial implementation process includes the development of an experimentally validated mathematical model, generation of a cohort of heterogeneous virtual patients, an assessment of stratification factors, and optimization of treatment strategy. We show the detailed process through application to melanoma treatment, using a combination therapy of chemotherapy and an AKT inhibitor, which was recently tested in a phase 1 clinical trial. We developed a mathematical model, composed of ordinary differential equations, based on experimental data showing that such therapies differentially induce autophagy in melanoma cells. Model parameters were estimated using an optimization algorithm that minimizes differences between predicted cell populations and experimentally measured cell numbers. The calibrated model was validated by comparing predicted cell populations with experimentally measured melanoma cell populations in twelve different treatment scheduling conditions. By using this validated model as the foundation for a genetic algorithm, we generated a cohort of virtual patients that mimics the heterogeneous combination therapy responses observed in a companion clinical trial. Sensitivity analysis of this cohort defined parameters that discriminated virtual patients having more favorable versus less favorable outcomes. Finally, the model predicts optimal therapeutic approaches across all virtual patients.

**One Sentence Summary:** We propose a computational framework to implement phase *i* trials (virtual/imaginary yet informed clinical trials) in cancer, using an experimentally calibrated mathematical model of melanoma combination therapy, that can readily capture observed heterogeneous clinical outcomes and be used to optimize future clinical trial design.

## Introduction

Significant advances have been made in understanding mechanisms that provoke tumor initiation and progression, and often this knowledge has been translated into the development of targeted agents that selectively disable the mutated, activated and/or overexpressed oncoproteins manifest in tumor cells (1). Most of these targeted agents have been tested in clinical trials either alone or in combination with other treatments (2), and though some are clinically effective (*e.g.,* small molecule *BRAF* kinase inhibitors (3)), the majority are not (4-6) despite the fact that such agents have potent activity in preclinical cancer cell and animal model studies. The leading cause of failure tends to be lack of efficacy, in part due to lack of robust predictive models that consider patient heterogeneity, and poorly designed clinical trials (6-9). This inconsistency is also partly due to difficulties in predicting the long-term effectiveness of a cancer therapy using time-limited *in vitro* (typically < 1 month) or *in vivo* (often < 3 months) models systems.

We reasoned that an appropriately defined and parameterized mathematical model, based on observations in cell and animal studies and clinical trials, might reveal insights regarding the design of improved and informed therapeutic approaches for treating cancer patients. To test this notion we used a mathematical model of melanoma that was generated based on data from *in vitro* experiments and a distribution of treatment responses (Fig 1A) from a comparable clinical trial. More specifically, the recently completed multi-arm phase 1 trial of the MK2206 AKT inhibitor (AKTi) in combination with standard chemotherapy (chemo) with advanced solid tumors, including melanomas (10) was used for these studies. To investigate potential mechanisms of treatment efficacy, a mathematical model comprised of a system of ordinary differential equations was developed to describe the dynamics of melanoma cells exposed to four treatment conditions, no treatment, chemo, AKTi and combination of chemo and AKTi. Cell culture experiments were then used to parameterize the model. The calibrated model was further validated using results from an extensive series of cell culture experiments that consider twelve different drug combinations and timings. This validated model was then used to predict the long-term effects of the twelve treatments on melanoma cells, which revealed that all treatments eventually fail, but do so at significantly different rates.

**Figure 1.**
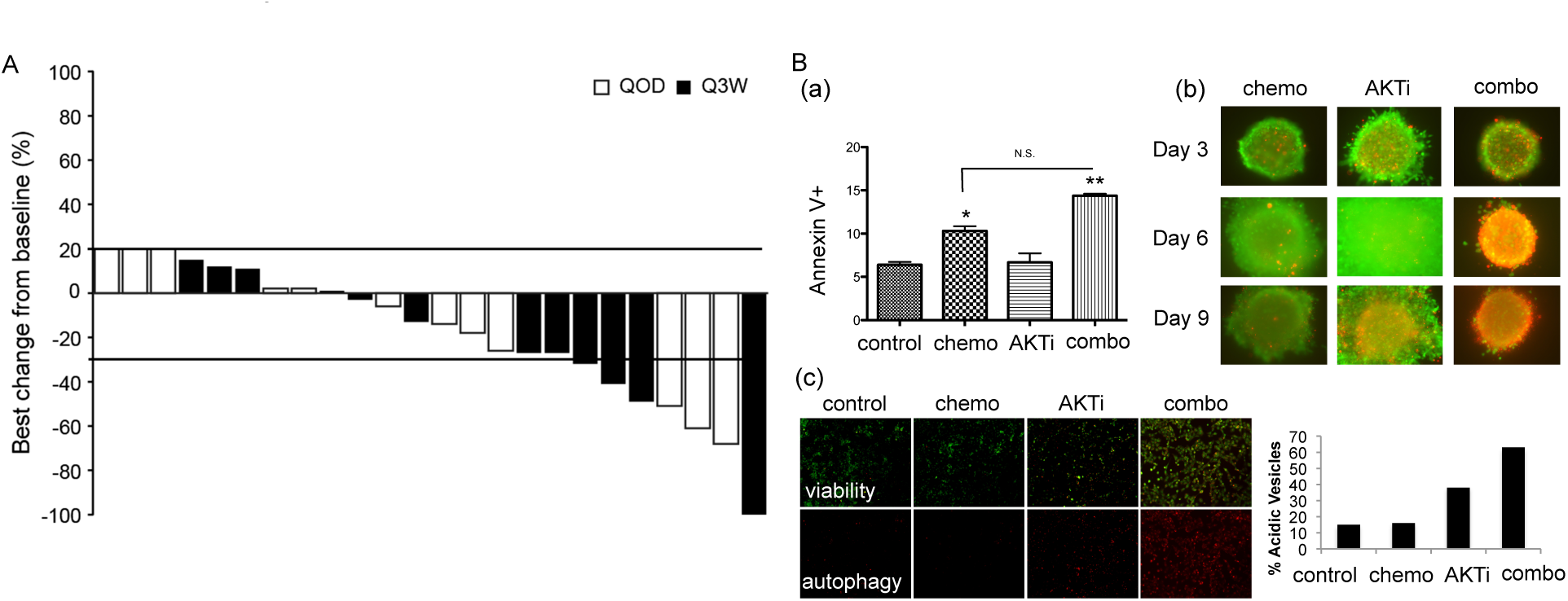
Acquisition of historic clinical and biological data. A, typical clinical data obtained from a clinical trial. Waterfall plots of the best responses in a clinical trial (10). The percentage change from baseline for the best result is shown here. Adapted from Molife et al (10), published in the Journal of Hematology & Oncology, copyright 2014 BioMed Central. B, biological data that may explain the underlying mechanisms in treatment responses. (a) Assessment of cell death (WM3918 cells, annexin-V) after 72 hr treatment with chemo (carboplatin and paclitaxel), AKTi (MK2206), and the combination of chemo and AKTi. (b) The measurement of cell death (WM3918) in a panel of 3D collagen-implemented spheroids. Cells were treated with AKTi and chemo (carboplatin and paclitaxel) for 3, 6, and 9 days. Green, viable cells; red, dead cells. Magnification 10x. (c) Treatment induces autophagy in cells. *Left,* fluorescence imaging of WM3918 cells treated as indicated for 72 hr and stained with acridine orange (AO). *Orange:* aggregated AO, *green:* diffuse AO. *Right,* Quantification of staining. Adapted from Rebecca et al (16) published in Pigment Cell & Melanoma Research, copyright 2014. John Wiley & Sons.

To investigate the long-term effects of therapy in a more clinically relevant setting, we varied model parameters to generate virtual patients that had a heterogeneous mix of responses similar to typical clinical trial outcomes (Fig. 1A and (10)). We employed a heuristic search algorithm (genetic algorithm (GA)) to generate a diverse virtual patient cohort consisting of over 3,000 patients. Treatment responses for this patient cohort were simulated and optimized, and the schedules of AKTi therapy were combined with chemotherapy. This strategy allowed implementation of a “virtual clinical trial” (phase *i* trial), where the model guides optimal treatment strategies for selected patient cohorts (11). Similar virtual clinical trials have been made to simulate clinical trials of cardiovascular disease, hypertension, diabetes (www.entelos.com), and acute inflammatory diseases (12). There have also been some previous studies that employed modeling approaches to predict outcomes of clinical trials (13, 14). Statistical approaches based on clinical drug metabolism (*e.g.,* dose-concentration relationships) have also been developed to design virtual clinical trials (reviewed in (15)) that detect significant differences between treatments (for example, placebo *vs.* treatment). Here we rather sought to translate biological findings coming from *in vitro* and clinical studies, using melanoma combination therapy, into a phase *i* trial - with the goal that this approach could be easily adapted to other cancers.

## Results

### Responses of melanoma treated with combination therapy

We reported unexpectedly long-term responses (of up to 15 months) to the combination therapy of chemo and AKTi in two *BRAF*-wild type melanoma patients in the trial (NCT00848718) (16). Although little was known regarding why this combination therapy was successful, we reasoned this reflected differential effects on inducing autophagy that were observed in Figure 1B. Autophagy represents a cancer cell-intrinsic mechanism of resistance that allows cells to survive times of drug-induced stress (17, 18) or, if uncontrolled, can deplete key cellular components and provoke tumor cell death (19, 20). Differential autophagic responses to the chemo plus AKTi combination and accompanying effects on tumor cell growth and survival were manifest in melanoma cells (Fig. 1B) (16).

### Mathematical model development

Motivated by these experimental results, we formulated a mathematical model comprised of three phenotypic compartments (Fig. 2A(a)), a non-autophagy compartment (N) and two autophagy compartments (P and Q). We divided the autophagy compartment into two, physiological autophagy (P) and quiescent autophagy (Q) compartments, based on studies showing that some cells where autophagy is manifest continue to maintain normal cell homeostasis whereas others did not (21-24). Figure 2A shows the interactions between the N, P and Q compartments within the four different environments: (a) no-treatment, (b) chemo, (c) AKTi, and (d) combination therapy (chemo+AKTi) (see the Materials and Methods section for a detailed description of model development).

**Figure 2.**
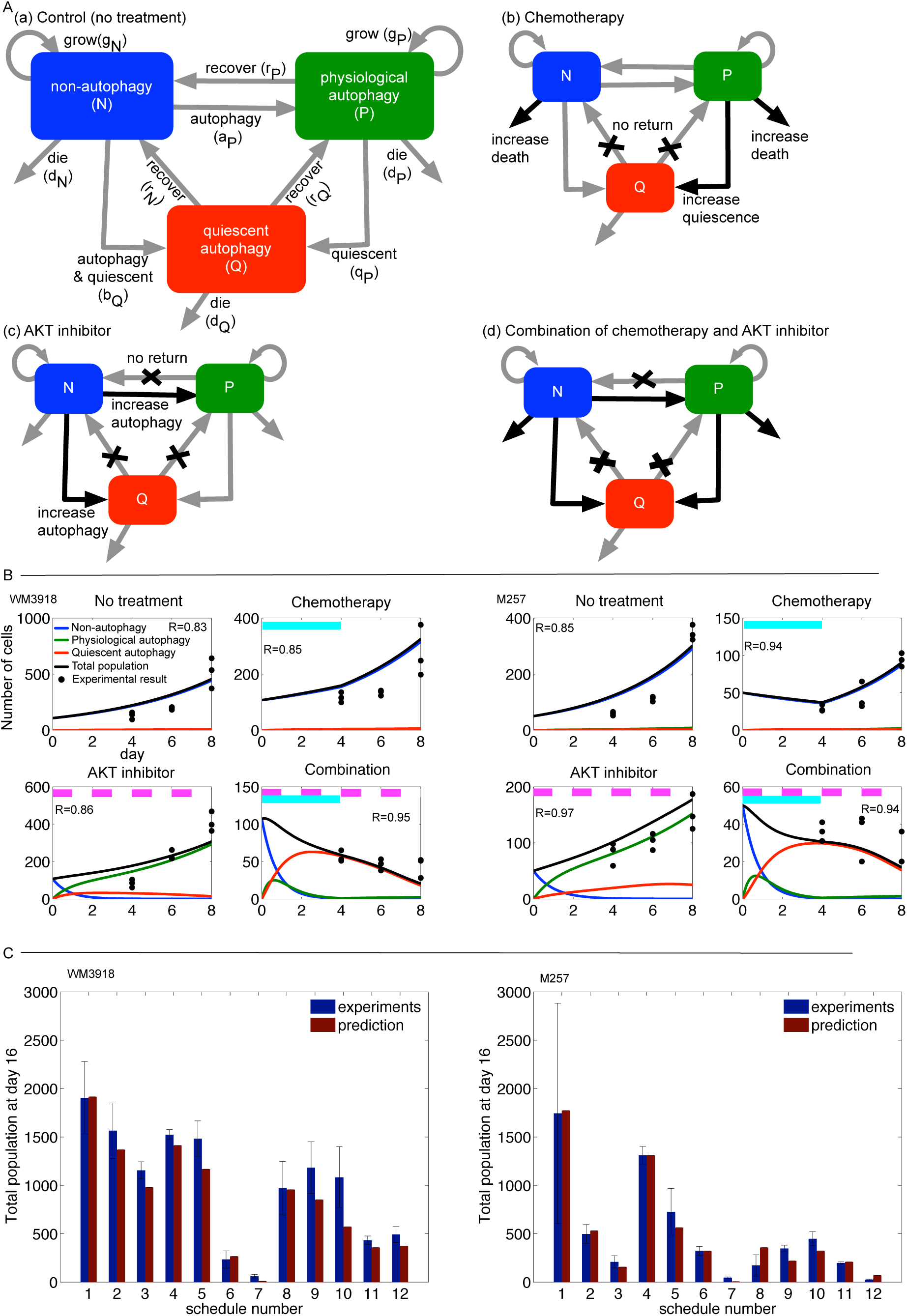
Mathematical model development and validation. A, Mathematical model development. (a) Schematic of a compartmental model composed of three compartments, non-autophagy (*blue*), physiological autophagy (*green*), and quiescent autophagy (*red*). (b) Schematic that incorporates the responses to chemo. *Black arrows,* increased rates during chemo; *black crosses,* the removal of the interaction during therapy. (c) AKTi increases the transition rate from non-autophagy to both autophagy phenotypes (*black arrows*) and blocks the transition from the autophagy to the non-autophagy phenotype (*black crosses*). (d) Schematic of the combination of chemo and AKTi. B, Model calibration. Predicted non-autophagy (*blue*), physiological autophagy (*green*), quiescent autophagy (*red*), total population (*black line*), and experimental cell counts (*black dots*) on days 4, 6 and 8, under four conditions: no treatment; chemo (day 0-4, *cyan bars*), AKTi (days 0, 2, 4 and 6, *pink bars*) and combination therapy (chemo on days 0-4 and AKTi on days 0, 2, 4 and 6) in cell lines WM 3918 (*left*) and M257 (*right*) melanoma cells. R-values are reported for each case. C, Validation. In a total of 12 different conditions (Fig. S1), each of the predicted total populations (*red bars*) are compared with of the each experimental results (*blue bars*).

To calibrate model parameters, two melanoma cell lines (M257 and WM3918) were treated with the four different conditions for 8 days and the results were compared with the model predictions. Using an optimization technique to minimize the difference between predicted and actual growth curves (see Materials and Methods) we obtained two suites of parameters that produced good fits (R ≥ 0.8, for all cases) for the growth of each cell line (Fig. 2B). Chemo reduced growth rates of both WM3918 and M257 (Fig. 2B, Chemo panels) and continuous application of AKTi increased the proportion of the non-autophagy (N) to physiological autophagy (P) phenotype (Fig. 2B, AKTi panels, increasing green lines). The combination of AKTi and chemo generally induced the quiescent autophagy (Q) phenotype (Fig. 2B, combination panels, increase of red lines), which arose following the rapid transition of a non-autophagy to physiological autophagy phenotype (Fig. 2B, combination panels, a sharp increase in the green line < day 1). The number of cells in the total cell population treated with the combination therapy continued to decrease as a result of cell death of the Q phenotype (Fig. 2B, combination panels, decreasing red lines).

### Model prediction and validation with preclinical data

The calibrated model was used to predict the effects of twelve treatment schedules that differed in the timing and order of chemo, AKTi and combination therapy across a 16-day period (Fig. S1, Materials and Methods). The expected treatment responses are summarized in Fig. 2C (red bars). One application of chemo decreased the tumor cell population by 30-65% (#2) and two applications reduced the population by 50-90% (#3). One application of AKTi had limited impact on tumor cell growth (approximately 20% reduction from #1 to #4). Continuous application of the AKTi reduced the tumor cell population size by 40-70% (#5). The M257 melanoma cell line was more sensitive to both chemo and AKTi than WM3918 melanoma cells. Combination therapies were substantially more effective than mono-therapies. In general, concurrent therapies with chemo and AKTi (#6 and #7) were more successful than all sequential therapies (#8-#12). Concurrent therapy #6 reduced the tumor cell population size by up to 90%, and concurrent therapy #7 nearly eradicated all of the tumor cells (Fig. 2C, #7, nearly invisible red bars). Sequential therapy decreased the tumor cell population size by 50-90% (Fig 2C #8-#12).

To validate these predictions, an extensive series of *in vitro* experiments were performed and the numbers of viable tumor cells were quantified on day 16 (Fig. S1B) using the Matlab Image Analysis toolbox. We then compared the total experimental melanoma cell numbers with those predicted by the models (Fig. 2C, blue *vs.* red bars). In general, the predictions matched well with the experimental results, as all predicted values (red bars) were within one standard error deviation from the mean value of the experimental results (blue bars). Thus, this mathematical model accurately describes and predicts treatment outcomes obtained *in vitro.*

### Long-term response of treatments

Having established a successfully validated model, the longer-term effects of the therapies were assessed. Surprisingly, the best proposed strategy (#7) failed by day 40 (WM3918 cells) or day 50 (M257 cells), where the physiological autophagy phenotype developed resistance to the therapy population (green lines, Fig. S2). This finding highlights a key shortcoming of *in vitro* experiments; *i.e.,* the limited timescale. To test this prediction long-term cell culture studies were performed. Perhaps unsurprisingly, as the model predicted, a 30-day treatment of strategy (#7) failed to eradicate some of the melanoma cell lines in colony formation assay experiments (16).

### Phase *i* trial: virtual cohort generation

Under the assumption that our underlying model mechanism is relevant to the clinic, and there is some evidence to support this (25), it was important to advance such analyses beyond homogeneous cell lines to the heterogeneous patient population. More specifically, in order to predict long-term treatment responses in a more clinically relevant scenario, treatments for model parameters that are more representative of real patients were simulated. The same treatment schedules as in the clinical trial (10) were used, with five cycles of first-day chemo in a 21-day cycle, and two different treatment arms of AKTi schedules (Q3W, QOD). In the first AKTi schedule (arm 1: Q3W), the drug is administered on the first day of the 21-day cycle for five cycles, and thereafter is given on the first day of every week (weekly maintenance therapy). In the second schedule (arm 2: QOD), the drug is administered on days 1, 3, 5 and 7 of the 21-day cycle, and is then given weekly as maintenance therapy.

To generate virtual patient cohorts that exhibit the diversity of responses observed in the clinic, adaptive heuristic search Gentetic Algorithms (GA, built in Matlab) were used, which are based on the evolutionary principle of natural selection (26, 27). A solution (*P* = [*g_N_, g_P_, d_Q_, d_τ_, r_p_, r_N_, b_p_, c_p_, c_Q_, a_N_, c_N_*]) was searched that minimized our objective function, 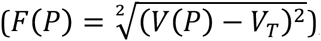, using the difference between tumor volume with a parameter (*V*(*P*) and the target tumor volume (*V_T_*). The *V_T_* was randomly selected from three categories (*V_C_, V_P_, V_S_*) based on response criteria (28), where subscript *C* denotes complete response (CR; tumor diameter < 1 mm), subscript *P* denotes partial response (PR; at least 30% reduction in tumor diameter) and subscript S denotes stable disease (SD; up to 20% increase in tumor diameter). Our GA found 1293 sets of parameters in the case of arm 1 (P_1_) and found 2098 sets of parameters for arm 2 (P_2_).

### Phase *i* trial: Stratifying treatment outcomes via clinically measurable parameters

To characterize this virtual cohort, it was first divided into four sub-cohorts (C_1_-C_4_, Fig. 3A(a)) according to the clinically measurable variables *c_N_*, chemo-sensitivity (29, 30), and *a_N_*), autophagy marker staining (31, 32). The hope being that this division might stratify best and worst responders. We compared a distribution of best tumor volume change responses of the 24 patients in the trial with that of 500 virtual patients sampled from our virtual cohort (3391 virtual patients) (Fig. 3A(b)). To test if two distributions are from the same continuous distribution, we performed a two-sample Kolmogorov-Smirnov test (Matlab statistics toolbox) (33). The test determined that we can not reject the null hypothesis at 5% significance (H = 0, P > 0.05, test statistic = 0.14). In addition, empirical cumulative distribution functions of the two overlap each other (Fig. 3A(b)). This implies that the best tumor volume responses of these 500 virtual patients can recapitulate real patients’ tumor volume response well. To validate the agreement further, we used Lin’s concordance correlation coefficient (ρ) (34, 35). We repeatedly sampled 24 virtual patients with an approximated probability density function of best tumor volume changes of real patients (Fig.1A) 1000 times. For each sample, we calculated ρ of two measurements, best tumor volume changes of both real patients and the sample, using an R package (epiR)(36) (Fig. S3). The resulting distribution of ρ shows that the two measurements are highly concordant (Fig. S3, ρ > 0.8).

**Figure 3.**
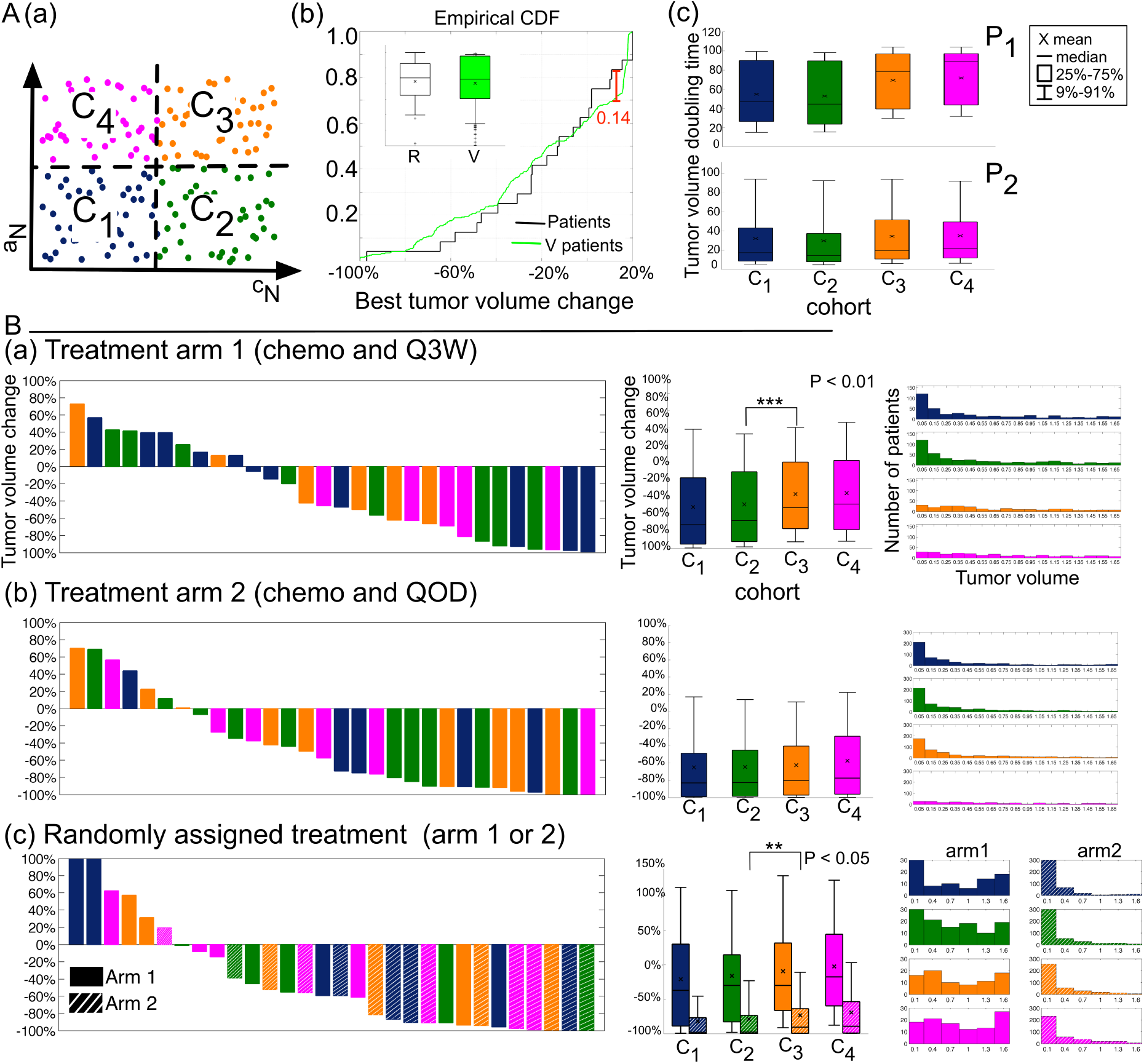
Virtual cohort generation and clinical trial simulation. A, Virtual cohort generation. (a) Graphical representation of virtual patients partitioned based on the cell death rate of the non-autophagy tumor population under chemo (*c_N_*) and the induction of the autophagy from non-autophagy to physiological autophagy populations following exposure to AKTi (*a_N_*). Both rates were low in patients in sub-cohort C_1_ (*dark blue dots*). *c_N_* was low but *a_N_* was high in C_2_ (*dark green*). Both rates were high in patients in C_3_ (*orange*). The tumor cells in patients in C_4_ (*pink*) had a higher rate of *a_N_* but a lower rate of *c_N_*. (b) Empirical cumulative distribution functions of patients (black line) and virtual patients (green line). The maximum difference between two distributions is 0.14. Inlet: Box-Whisker plots of patients (left) and sampled 500 virtual patients (right). x: mean, -: median, box: 25% - 75%, upper and lower horizontal bar (-): 91-9%.(c) Mean doubling time of untreated tumor volume in virtual patients from group P_1_ (top), P_2_ (bottom). *Dark blue*: C_1_; *dark green*, C_2_; *orange*, C_3_; and *pink,* C_4_. B, Simulation of 6-month treatments in all of the virtual patients. (a) Waterfall plots of final tumor volumes in 30 randomly selected patients from P_1_ treated with arm 1. Box whisker plot of the expected 6-month treatment outcomes in each sub-cohort C_1-4_ (×: mean, -: median, box: 25% - 75%, upper and lower horizontal bar (-): 91-9%). **, P < 0.05; ***, P < 0.01. Histogram shows expected tumor volumes at the final day of treatment (day 180). (b) Waterfall plots of final tumor volume in 30 randomly selected patients from P_2_ treated with arm 2. Box whisker plot of the expected 6-month treatment outcomes in each sub-cohort C_1-4_ Histogram of expected tumor volumes. (c) Waterfall plots of tumor volume response in 30 randomly selected patients from both P_1_ and P_2_ treated with either arm 1 or arm 2 (randomly selected treatment). Box whisker plot of the expected 6-month treatment outcomes in each sub-cohort C_1-_C_4_. Histogram of expected tumor volumes (day 180).

The native doubling time (*i.e.,* without any treatment) of each sub-cohort (C_1_-C_4_) in each patient group (P_1_ and P_2_) was compared assuming exponential tumor growth, which is reasonable for metastatic disease, and the exponent of the best-fit curve was determined. Doubling times were in the range of 7 to 110 days (Fig. 3A(c) top P_1_, Fig. 3A(c) bottom for P_2_), which were similar to values observed in patients with cutaneous melanoma (37). Overall, tumors in the P_2_ group grew much faster than those in P_1_ (mean of ~30 days *vs.* mean of ~60 days, respectively). In P_1_, tumors in C_1_ (dark blue) and C_2_ (dark green) that had slightly increased autophagy following AKTi treatment grew faster than those in C_3_ (orange) and C_4_ (pink) that had highly increased levels of autophagy (50 days *vs.* 70 days). In contrast, there were no significant differences in the doubling times between the sub-cohorts in P_2_.

Phase *i* combination therapy was simulated for 6 months with the two arms to predict treatment responses in the virtual cohorts. Patients in P_1_ treated with arm 1 had diverse responses to the therapy, where CR, PR and SD were observed in 30 randomly selected subjects (Fig. 3B(a), waterfall plot). For example, some patients in C_1_ were expected to be complete responders ( CR, dark blue bars of nearly -100%), while others in the same cohort were expected to be in SD (dark blue bars of 40-60%). When comparing the expected tumor volume changes in each cohort (Fig 3B(a), middle boxplot), there were no significant differences in outcomes between C_1_ and C_2_ or between C_3_ and C_4_. However, the C_2_ and C_3_ cohorts showed a significant difference (P <0.01, Student’s *t*-test). Thus, one of the parameters, specifically the physiological autophagy phenotype (*a_N_*), defines outcome. The histogram of all subjects in each cohort also showed a shift in treatment responses from the cohort C_1-2_ (skewed toward responsive) to C_3-4_ (uniformly distributed tumor volumes).

Similarly, subjects in P_2_ treated with arm 2 also showed diverse responses to therapy (Fig. 3B(b)). However, unlike subjects in P_1_ (Fig. 3B(a)), none of the metrics determined outcomes of subjects in P_2_, as the mean tumor volume responses in the four cohorts did not show significant differences (Fig. 3B(b), boxplot). Accordingly, the histograms only showed a slight shift of responses when C_1-3_ was compared to C_4_ (Fig. 3B(b)).

We also randomly assigned a treatment arm (either arm 1 or arm 2) to all virtual patients in both P_1_ and P_2_. Tumor volume was checked at the beginning of each cycle (every 21 days), and subjects were removed from treatment if the tumor volume had more than doubled; this resulted in the removal of a total of 1473 subjects. Random assignment of the treatment arms produced even greater variations in expected outcomes (Fig. 3B(c)), but might more accurately reflect how an actual population responds. Specifically, the expected tumor volume changes showed a large variation, where the 9-99% bar in the boxplot increased in all sub-cohorts, and in both arms. The physiological autophagy phenotype (*a_N_*) distinguished outcomes between C_2_ and C_3_ only when patients were treated with arm 2 (Fig. 3B(c), boxplot, P <0.05). There was no shift in the distribution of responses between the cohorts (Fig. 3B(c), histograms). Interestingly, the outcomes were almost uniformly distributed in patients treated with arm 1, while the distributions of patients treated with arm 2 were skewed toward smaller final tumor volumes (more responsive to the therapy). Thus, the physiological autophagy phenotype defines outcomes in diverse virtual cohorts.

### Phase *i* trial: Assessing stratification factors by sensitivity analysis

Rather than focusing solely on the clinically quantifiable parameters, the nine remaining model parameters were also assessed to identify sub-cohorts that are most or least likely to benefit from the combination therapy. To identify the key parameters a sensitivity analysis of the effect of each parameter on tumor volume after 6 months of treatment was performed. After dividing the patient group into CR, PR, and SD (based on their tumor volume at 6 months), each parameter value from each group was compared and the Student’s *t*-test was used determine if the mean values of a parameter differed between two groups (CR *vs.* PR and PR *vs.* SD) (Fig. S4-S6). In patients in P_1_, the two autophagy-associated phenotypes, physiological and quiescent (*a_N_, b_P_*), were significantly different between CR and PR and between PR and SD (Fig. S4). Compared with the previous partition (Fig. 3A), the new partitioning of P_1_ according to the value of *a_N_* and *b_P_* (Fig. 4A,) discriminated the expected treatment outcomes more clearly. Unlike the previous partition result (Fig. 3B(a)), each sub-cohort (C_1-4_) now showed significantly different mean response to therapy (Fig. 4A, boxplot). The new sub-cohort C_1_ (cyan) is most likely to benefit from the treatment, while the likelihood decreased gradually in C_2_ (blue), C_3_ (yellow), and C_4_ (red). There is also a clear shift in the distribution of treatment response from C_1_ to C_4_ (Fig. 4A, waterfall plot).

**Figure 4.**
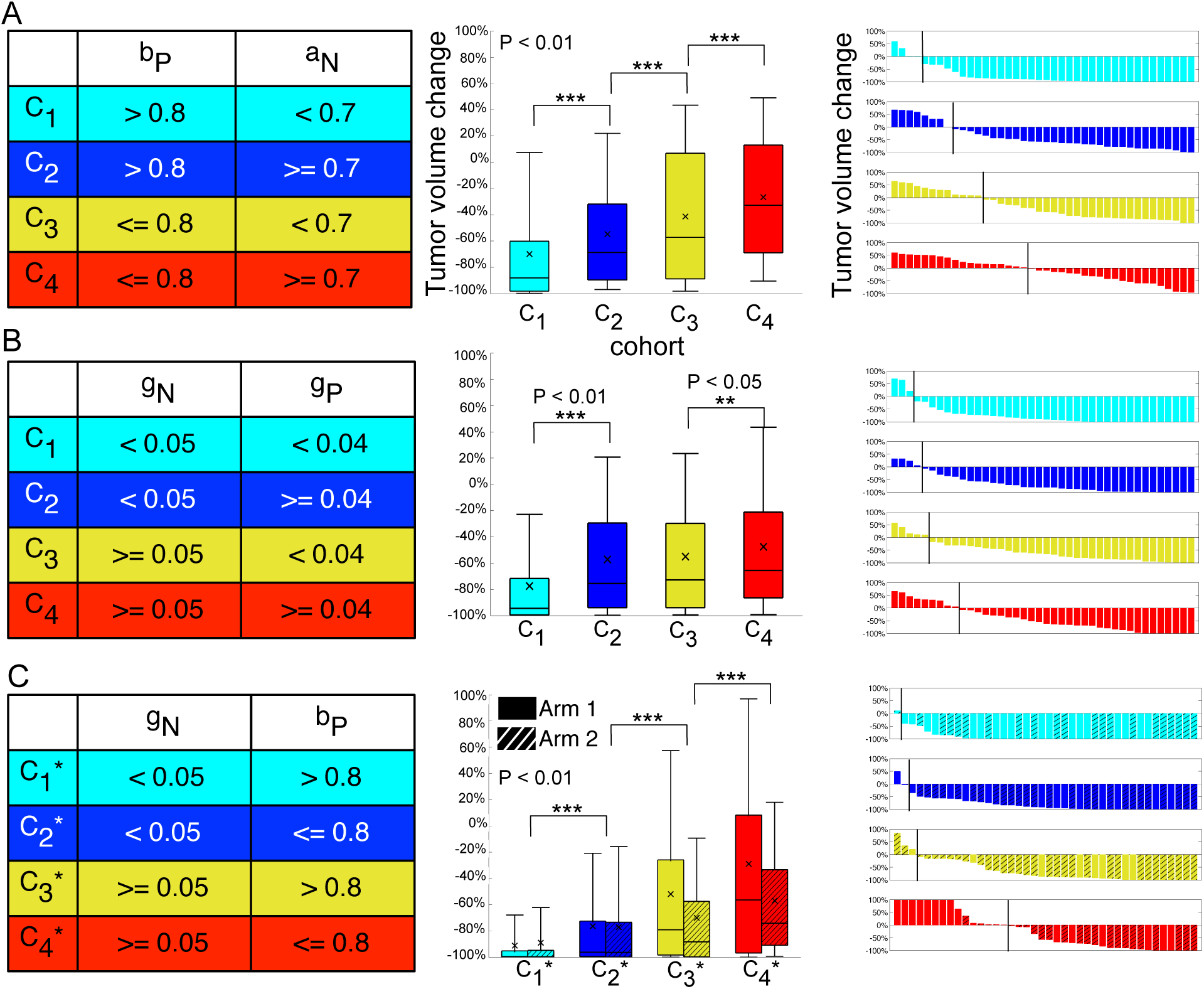
Potential predictive factors of the therapy to virtual patients. A, Re-partitioning of P_1_ based on the rate of autophagy induction in the transition from the non-autophagy to physiological autophagy phenotypes under AKTi (*a_N_*), and the rate of transition from the non-autophagy to quiescent autophagy phenotypes (*b_p_*). Box whisker plot of the expected 6-month treatment outcomes in the new sub-cohorts C_1-_C_4_ of P_1_ (x: mean, -: median, box: 25% - 75%, upper and lower horizontal bar (-): 91-9%). *Cyan,* C_1_; *blue,* C_2_; *yellow,* C_3_; and *red,* C_4_. **, **P** < 0.05; ***, **P** < 0.01. Waterfall plots of final tumor volumes in 40 randomly selected patients from P_1_. The vertical line indicates no change in volume (0%) after therapy. B, Re-partitioning of **P**_2_ based on two growth rates, those of non-autophagy (*g_N_*) and physiological autophagy phenotype cells (*g_p_*). Box whisker plot of the expected 6-month treatment outcomes in the new sub-cohorts C_1-_C_4_ of P_2_. Waterfall plots of final tumor volumes in 40 randomly selected patients from P_2_. C, Re-partitioning of all virtual patients based on the growth rate of non-autophagy tumor cells (*g_N_*) and the transition rate from non-autophagy to quiescent autophagy phenotype (*b_p_*). Box whisker plot of the expected 6-month treatment outcomes in the new sub-cohorts C_1_^*^-C_4_^*^. Waterfall plots of the final tumor volumes in 40 randomly selected 40 patients treated with either arm 1 or arm 2 (random assignment).

Similarly, our sensitivity analysis on P_2_ (Fig. S5) revealed another key parameter set (*g_N_, g_P_*), the growth rates of both the non-autophagy and physiological autophagy phenotypes. P_2_ was divided based on the parameter values (C_1_-C_4_) (Fig. 4B, table). Unlike the previous partitioning (Fig. 3B(b)), these parameters successfully segregated patients according to the expected outcomes (Fig. 4B, boxplot). C_1_ was revealed as the best responding sub-cohort, where almost all tumor volumes in this group (99%) were reduced by at least 20% (Fig. 4B, cyan boxplot). The next best responders were both C_2_ and C_3_, where more than 75% of the cohorts showed at least a 30% reduction in tumor volume; the expected outcomes in C_2_ and C_3_ significantly differed from those in C_1_(P < 0.01). The expected outcomes in C_4_ were slightly worse than those in C_1_-C_3_, as the mean outcome was worse with some tumors from C_4_ having an increased tumor size of up to ~70% (Fig. 4B, red boxplot). The waterfall plots in Fig. 4B show inter-patient variation in tumor volume changes even within the same sub-cohort and shifting of the treatment response from C_1_ to C_4_.

With random treatment assignment of either arm 1 or arm 2 to all patients in both P_1_ and P_2_, as would be done in a real trial, our analysis identified the key parameters *g_N_* and *a_N_* as potential predictive factors of the treatment outcomes (Fig. S6). The new cohorts (C_1_^*^-C_4_^*^, Table in Fig. 4C) exhibit significantly different mean responses to therapy (Fig. 4C, boxplots). The new partition also showed a tendency of increasing tumor volumes from C_1_^*^ to C_4_^*^ in both arms (Fig. 4C, boxplots and waterfall plots), and defined the most sensitive sub-cohort (C_1_^*^), who are predicted to be the most likely to benefit from treatment using either arm 1 or arm 2. In summary, sensitivity analysis allows one to select potential predictive factors for each treatment strategy, which could be used as a guide for selecting specific therapies for a given patient.

### Phase *i* trial: Optimizing AKTi treatment to minimize toxicity for each sub-cohort

For the new virtual sub-cohorts (C_1_^*^-C_4_^*^) optimal therapy recommendations were derived using implicit filtering (38). Notably, the schedule of chemo was fixed, as it was in the clinical trial (10), with the goals of identifying the AKTi schedule that reduced the initial tumor volume by at least 30% after 6 months of therapy, and that used as few AKTi applications as possible to provide an effective and less toxic treatment strategy. Optimized schedules of AKTi are summarized in Table 1. For the patients in C_1_^*^, optimum therapy recommended 1 day of AKTi treatment on the first day of the 42-day cycle, a treatment holiday for five cycles and then from day 168, 1 day of treatment on the first day of a 7-day cycle (weekly maintenance). For the patients in C_2_^*^, the optimal schedule was to administer AKTi on the first and third days of a 21-day cycle for five cycles, followed by weekly maintenance. For the patients in C_3_^*^, optimization recommended that AKTi be taken on the first, third, and fifth days of each 19-day cycle for five cycles, followed by administration on the first day of each week from day 95. The optimal schedule for cohort C_4_^*^ is to administer AKTi on the first, third, and fifth days of each 19-day cycle for five cycles, followed by administration on the first and third days of a 9-day cycle from day 96.

**Table 1.**
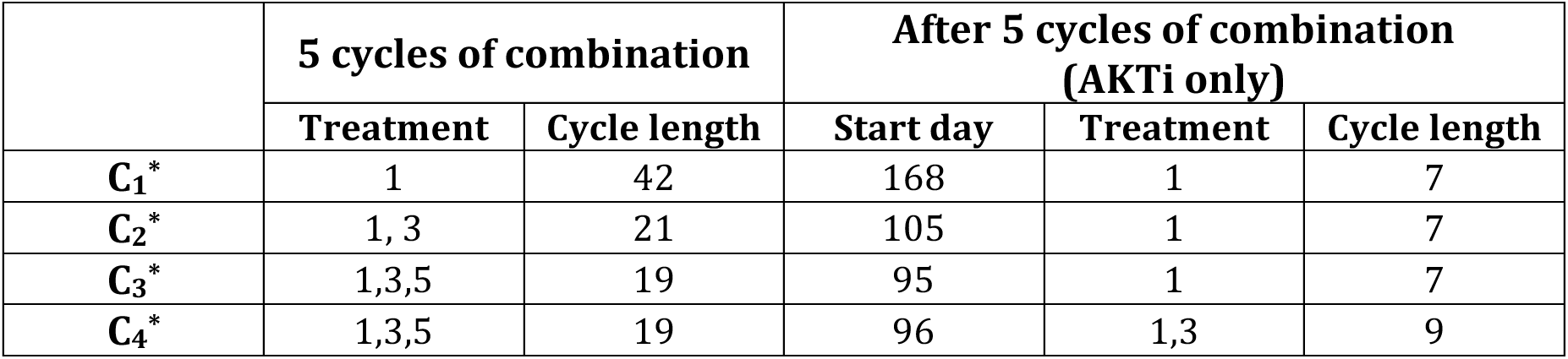
Optimized AKTi schedule for sub-cohort C_1-4_^*^ (unit: day)

To understand the relative impact of these alternate therapies we compared the 2-year survival probability of patients on the different therapies. The therapies included optimized therapy, treatment arm 1 and arm 2, and two mono-therapies of AKTi only (*i.e.,* AKTi treatment only from either arm 1 or arm 2), and chemo (Fig. 5). The initial number of tumor cells was set at one billion and the melanoma was considered fatal when the number of tumor cells reached 10^13^ cells. We also compared the cumulative drug concentration of each therapy to determine the toxicity of the treatment at two time points, immediately after the five-cycles of combination therapy of AKTi with chemo (105 days) and at the completion of therapy (2 years) (see Materials and Methods for calculations of plasma concentration).

As expected, the probability of 2-year survival decreased after 6 months if the patients in the cohort C_1_^*^ were not treated with any therapy (Fig. 5A, black line). Survival of the chemo-alone C_1_^*^ cohort had only minimal improvement (Fig. 5A, yellow line). In contrast if the C_1_^*^ patients were treated with any type of AKTi, their probability for 2 year survival remained over 0.9 (Fig. 5A, red, pink, blue, cyan and green lines), which was expected as this cohort is the most sensitive (Fig. 4C, cyan boxplots, tumor volume at 6 months < -60%). Regarding toxicity, the optimal therapy was the least toxic to patients, as the cumulative drug concentration was minimal (Fig. 5A, green bars), except the C_1_^*^ chemo cohort (Fig. 5A, nearly invisible yellow bar).

**Figure 5.**
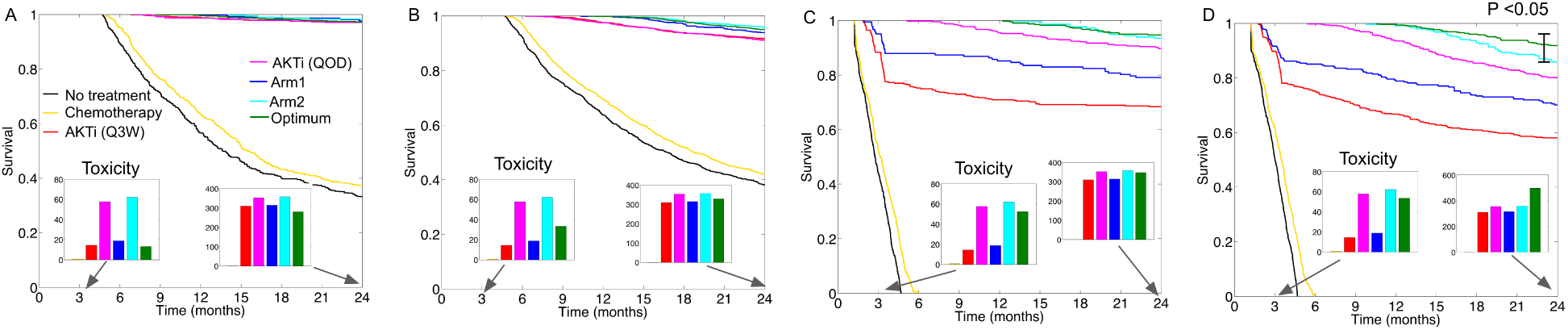
Treatment optimization and predicted 5-year survival analysis. Kaplan-Meier curves in patients from cohort (A: C_1_^*^, B: C_2_^*^, C: C_3_^*^, D: C_4_^*^) when eight different treatments were applied for 2 years, no treatment (*black*), five cycles of chemo only (*yellow*), AKTi monotherapy following the Q3W schedule (*red*), AKTi monotherapy following the QOD schedule (*pink*), combination therapy with arm 1 (*blue*) or arm 2 (*cyan*), and the optimum therapy (*green*). Inlet bars, sum of cumulative drug (both AKTi and chemo) concentrations at day 105 (first bar graph, same color scheme as in the survival curves) and year 2 (second bar graph).

The probability of 2-year survival of patients in cohort C_2_^*^ also decreased after 6 months if patients were not treated with any therapy (Fig. 5B, black line) and chemo again slightly increased the probability of survival (Fig. 5B, yellow line). Also, like the cohort C_1_^*^, the probability of 2-year survival was relatively high (> 0.9) when patients were treated with any of the AKTi schedules (Fig. 5B). The combination AKTi with chemo and the optimized therapy slightly increased survival probability (Fig. 5B, cyan, blue, and green lines). Regarding toxicity, arm 2 (cyan) was the most toxic, followed by AKTi (QOD, pink), the optimized therapy (green bar), and arm 1 (blue bar), at both days 105 and 720 (2 years) (Fig. 5B bar graphs). Interestingly, the most optimized therapy is substantially less toxic than both arm 2 and AKTi (QOD) at day 105.

For patient cohort C_3_^*^, the survival probability decreased rapidly to zero by 6 months if the patients were untreated (Fig. 5C, black line) and chemo again had little-to-no. impact on survival (Fig. 5C, yellow line). The AKTi monotherapies and treatment arm 1 significantly improved patient survival (Fig. 5C, AKTi: pink and red lines and arm 1: blue line). However, the most optimized therapy and arm 2 clearly provided the best probability for survival, and again the optimized therapy was less toxic than arm 2 (Fig. 5C, green *vs.* cyan bars).

Finally, like the C_3_^*^ cohort, the probability of survival of C_4_^*^ patients decreased sharply if they were untreated or treated with chemo only (Fig. 5D, black and yellow line), and again the AKTi monotherapies improved survival (Fig. 5D, red line (Q3W), and pink line (QOD)). Both treatment arm 1 and arm 2 also increased the probability of survival (Fig. 5D, blue line: arm 1 and cyan line: arm 2). Notably, the optimized therapy significantly improved the probability of survival (Fig. 5D, green line) compared to arm 2 (P < 0.05), and was also less toxic than arm 2 at day 105 right after the five cycles of combination of chemo with AKTi, although it became more toxic at year 2 (Fig. 5D bar graphs).

## Discussion

The integrated approach applied herein shows that treatment-induced autophagy phenotypes is certainly one factor driving the long-term effects of treating patients with chemo in combination with AKTi. The mathematical model hypothesizes that two distinct states of autophagy exists (physiological and quiescent states), and indicates that improved patient outcomes are associated with the quiescent autophagy phenotype. The model also predicts that therapy drives the transition from the non-autophagy to the physiological autophagy phenotype, which provides a transient escape route from treatments. In contrast, the model indicates that a persistent quiescent autophagy state is detrimental to overall fitness and thus this represents a desired outcome of therapy.

Implementing a phase *i* trial allows one to translate models to a clinically relevant setting. The key components of such a trial are an experimentally calibrated mathematical model and a cohort of virtual patients that mirror responses observed in an actual clinical trial. When developing mathematical models to facilitate clinical decisions, an important constraint is the number of measurable parameters of the patient. As patient data are generally limiting, this inevitably leads to simpler models, where the data dictate model inputs and consequently, model outputs. The model presented herein is an excellent example, as it is constructed by integrating experimental findings for a given cancer (*e.g.,* melanoma) with the clinical reality. An extensive cohort having widely distributed parameters is another key element of the phase *i* trial, and the cohort should be representative of inter-patient variability, which drives the diversity of the response (*i.e.,* the spectrum of the response and resistance to a given therapy) that is observed in the clinic. Using a GA and employing response criteria used in the clinic, one can easily generate a relatively large cohort of diverse virtual patients who are treated using the same treatments given to patients in a clinical trial and statistically reproduce the same responses.

To stratify treatment outcomes according to clinically measurable variables, the cohort was divided based on chemo-sensitivity and the induction of autophagy in tumor samples. The rate of transition to the physiological autophagy phenotype was a predictive factor of treatment outcome for arm 1, where higher rates of physiological autophagy resulted in unfavorable outcomes. Interestingly in another melanoma study using different treatments, an increased autophagy response was associated with resistance to *BRAF* inhibitors (25). In treatment arm 2, neither of the two variables selected for more sensitive or resistant individuals. However, after randomly assigning treatment of either arm 1 or arm 2 to each virtual patient, the autophagy phenotype again discriminated sensitive *vs.* non-sensitive sub-cohorts. Interestingly, some of the good responders to arm 2 became non-responders when their treatment was switched to arm 1 (Fig. 3B(c), 100% dark blue bars on the far left), highlighting the importance of knowing the most appropriate patients to be assigned to each treatment arm.

Sensitivity analysis of the simulated cohort was superior at defining predictive factors in each treatment arm, which discriminated between CR, PR and SD. The ability of non-autophagic melanoma cells to become either physiological or quiescent autophagic determined patient outcomes for arm 1, whereas the growth rates of proliferating tumor cells (non-autophagy *vs.* autophagy phenotypes) were the determinants of outcomes for arm 2. Further, when randomly assigning to either arm 1 or arm 2 the transition to the quiescent autophagy phenotype and the growth rate of non-autophagy cells discriminated outcomes. Thus, key parameters for cases having more or less favorable outcomes can be defined for each treatment schedule, demonstrating the utility of phase *i* trials in aiding patient selection.

Optimizing the AKTi schedule for each cohort provided the most benefit. Notably, mathematically informed drug scheduling can positively impact overall outcome, including using a lower drug dose in some cohorts. Indeed, changing the temporal protocol influenced the dynamics of the system significantly. Interestingly, another melanoma study showed that using unconventional (discontinuous) dosing schedules of melanoma cells with *BRAF* inhibitors could prevent resistance (39). This idea is now being explored in a phase II clinical trial of *BRAF*-mutant melanoma patients (SWOG: 1320). We submit that the simulation of optimized schedules and comparing outcomes across virtual patients can assist clinical treatment planning to improve overall outcomes (10).

The underlining mechanisms, parameterization and validation of our model were based on data from a preclinical *in vitro* study. However, as demonstrated Leder and colleagues an integrated approach can be also achieved using *in vivo* preclinical studies (40). To bridge the divide between our *in vitro* study and the clinic the assumption was made that the same mechanisms of therapy resistance apply. Although there are a multitude of potential resistance mechanisms in patients, we consider one (autophagy) that we characterized using our integrated approach of mathematical modeling with *in vitro* experiments. We also assume that the variability in patients’ responses to treatment can be characterized by variability in this autophagy mechanism, although there are certainly more sources of variability including but not limited to the variation in patient age, size and the genetic composition of an individual tumor, and immune responses. Whilst it would certainly be possible to incorporate both additional resistance mechanisms and sources of variability into our model, and given appropriate experimental controls to calibrate and validate our model, ultimately the methodology would be the same, albeit with many more parameters to generate virtual cohorts. What is clear, regardless of the model complexity or cancer system, is that using a validated mathematical model (with pre-clinical data) to generate a virtual patient cohort (that matches the distribution of clinically observed outcomes) will allow us to carry out phase *i* trials that can be broadly applied to improve the safety and efficacy of future phase I-IV trials, as well as patient outcomes.

## Materials and Methods

### Mathematical modeling

Untreated melanoma cells proliferate (rate:*g_N_*), and following treatment can acquire either a physiological autophagy phenotype (transition rate:*a_p_*) or a quiescent autophagy phenotype (transition rate: *b_Q_*). Physiological autophagy cells grow (rate: *g_P_*) and can revert to non-autophagy cells (returning rate:*r_p_*), or enter a quiescent/senescent state (rate:*q_p_*). Tumor cells having the quiescent autophagy phenotype do not divide, yet such these cells can either reacquire a physiological autophagy phenotype (rate: *r_Q_*) or a non-autophagy phenotype (rate: *r_N_*) state. Cells in each compartment die at some rate (*d_N,P,Q_*). To model increased cell deaths on days 6-9 (Fig. 1B(b)), we included the delayed cell deaths of quiescent/senescent autophagy cells (*τ*).

The effects of chemo, AKTi and their combination (Fig. 2A(b)-(d)) were incorporated into the model. It was assumed that the drug reaches its maximum concentration immediately after administration and remains at that level until the beginning of the treatment break. Although the chemotherapeutic agents (paclitaxel and carboplatin) are detectable in patients for 24 hours, the half-life in serum is relatively short, in the range of 5.6 – 11.1 hr (41, 42); we assumed that the concentration of the chemotherapeutic agents were maintained for only 1 day after administration and became zero at the beginning of the treatment break. As the plasma concentration of AKTi (MK2206) is known to be constant for approximately 48 hr (with a long terminal elimination half-life of 40-100 hr) (43), we assumed that a 1-day application of AKTi to cells or patients corresponds to a 2-day application to cells or patients *in silico.* Notably, drug doses were not modulated in this study, where a fixed dose was assumed for both chemo and AKTi. As cell culture experiments showed that chemo triggered cell death with negligible effects on autophagy (Fig. 1B(a)), it was assumed that chemo only augmented cell death. Further, as chemo is effective only in proliferating melanoma populations, we assumed that the therapy increases the death rate of the two proliferating phenotypes, non-autophagy (*d_N_*) and physiological autophagy cells (*d_p_*)(Fig. 2B(b), black arrows). We also assumed that the frequency with which cells became quiescent/senescent (*q_p_*) increased with chemo (Fig. 2B(b), black lines). *In vitro* studies showed that while AKTi did not augment cell deaths or effectively inhibit melanoma cell growth (Fig. 1B(a)-(b) and (16)), it did induce autophagy (Fig. 1B(c) and (16)); thus, we assumed that AKTi increases the rate of transitioning to the autophagy phenotypes, *a_p_* and *b_Q_* (Fig. 2B(c), black arrows). As combination therapy does not augment cell death compared with chemo, nor significantly increase autophagy relative to AKTi, the combination of the two treatments was modeled by adding the effects of chemo and AKTi (Fig. 2B(d), black arrows and crosses). Finally, it was assumed that no cells with a given phenotype revert to their original states while any treatment is being applied. The schematic representation of this compartment model (Fig. 2A(a) to (d)) converts readily into a system of ordinary differential equations:

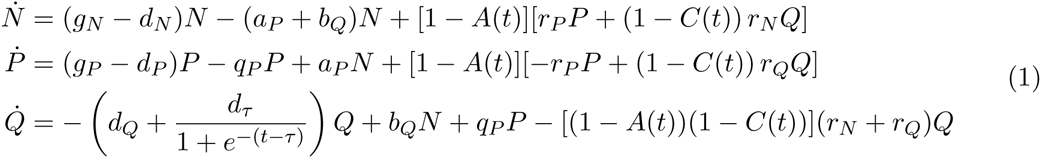

where *d_N_* = *d*_0_ + *c_N_C*(*t*), *d_P_* = *d*_0_ + *c_P_C*(*t*), *q_P_* = *q*_0_ + *c_Q_C*(*t*), *a_P_* = *a*_0_ + *a_N_A*(*t*), and *b_Q_* = *b*_0_ + *b_P_A*(*t*). In equation (1), *A*(*t*) and *C*(*t*) model time schedules of AKTi and chemo, respectively. To model the treatment schedules, we used Heaviside function *H* (*t* − *θ*) defined by

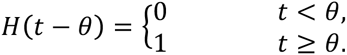

Assuming alternative application of the AKTi over a period of time at each cycle, the schedule of AKTi was modeled by 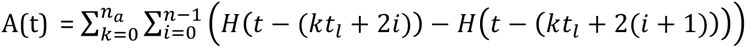 where *n* is the number of AKTi applications at each cycle, *n_a_* is the number of AKTi treatment cycles, and *t_l_* is the length of each AKTi treatment cycle. Similarly, the schedule of chemo was modeled by 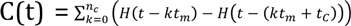, where *n_c_* is the number of chemo cycles, *t_m_* is the length of each chemo-cycle, and *t_c_* is the duration of chemo. The equation (1) was solved using a Matlab ODE solver (ode15s).

### Parameter estimation

Two melanoma cell lines (M257 and WM3918) were treated with chemo on the first four days (day 0-4) and with AKTi on every other day (days 0, 2, 4, and 6). We measured the number of melanoma cells at days 4, 6 and 8. We assumed that all of the initial tumor cell population was the non-autophagy phenotype (100% of non-autophagy cells on day 0). Both initial cell populations and the growth rate of non-autophagy cells (*g_N_*) are estimated by assuming an exponential growth of untreated cells and finding both the initial value and exponent of the best-fit curve. We used an optimization algorithm called implicit filtering (38), a steepest descent algorithm for problems with bound constraint, to determine the best remaining parameter set *H* (except *g_N_*) that minimized the difference between predicted number of cells (*N_P_*) and experimental result (*N_E_*) in four conditions: no treatment (*n*); chemo (*c*); AKTi (*a*); and combination (*m*). The mathematical definition of our problem is:

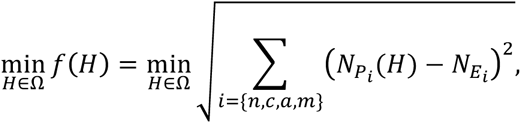

where the goal is to minimize the objective function *f* subject to the condition that 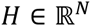 is in the feasible region:

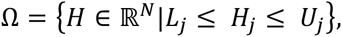

where and *L_j_* and *U_j_* are the upper and lower bound on the *j*th component *H_j_* of the vector *H*. The estimated parameters for each cell line are summarized in Table S1.

### Synopsis of 12 different schedules over 16 days tested to validate the model

The schedules (Fig. S1) included: #1, no-treatment; #2, two types of mono-therapies with chemo only (one time chemo for the first four days, days 0-4); #3, two chemos for both the first four days (days 0-4) and the last four days (days 12-16); #4, AKTi mono-therapy (one time AKTi for the first four days (days 0-4), #5: AKTi mono-therapy, alternative day administration of AKTi.

We considered both concurrent and sequential combination therapies. The two concurrent combination therapy schedules are #6 and #7, where #6 was one time treatment of concurrent therapy for the first four days (day 0-4), and where #7 was one time treatment of chemo for the first 4 days (days 0-4) and alternative administration of AKTi.

The four single agent sequential therapy schedules (#8-#12) included different ordering of chemo and AKTi. Some schedules set chemo treatment first, where #8 was chemo from day 0 to day 4 and then AKTi from day 0 to day 4, whereas #9 was chemo on day 0-4 and then alternating administration of AKTi. The other schedule administrated AKTi first, where #10 was AKTi first for day 0-4 and then chemo day 0-4. Finally, there were alternative applications of chemo and AKTi twice, where #11 was chemo first for the first four days, followed by AKTi treatment for 4 days, and where #12 was AKTi for the first four days, followed by 4-day application of chemo.

### Optimization of AKTi schedules

In the model AKTi was initially administered with chemo for five cycles and then administered without chemo (AKTi maintenance), as in the clinical trial. For each AKTi application, we assumed a 1-day application time. The scheduling parameter *S* consists of 5 variables, number of 1-day applications (*N_c_*) at each cycle with chemo, length of treatment holiday at each cycle (*L_c_*), starting day of AKTi maintenance (*M_s_*), number of applications (*N_s_*) in AKTi maintenance phase and the length of treatment holiday (*L_s_*) each week since the stating day *A_s_*. Our objective function to identify an optimal schedule of the AKTi is 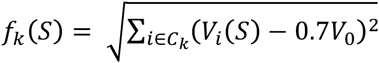, where k (=1,2,3,4) denotes each cohort C_1-4_^*^and *V_0_* is the initial number of tumor cells (10^9^ cells). We employed the implicit filtering method to find the optimal S for each cohort.

### Toxicity treatment: cumulative drug concentration of AKTi and chemo

The plasma concentration of AKTi was determined by solving a differential equation 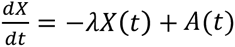, where *λ* is a decay rate of AKTi and *A*(*t*) (defined in the above Mathematical modeling section) denotes the application of AKTi. We assumed that the decay rate of AKTi is 0.35 per day based on pharmacokinetic study of AKTi in patients (43). Similarly, plasma concentration of chemo was obtained by solving a differential equation 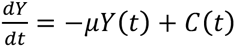, where *μ* (= 0.7 per day) is a decay rate of chemo and *C*(*t*) denotes the application chemo (see Mathematical modeling section).The cumulative drug concentration at time *T* is obtained by adding the areas under the curve of *X*(*t*) (AKTi) and *Y*(*t*)(chemo) from 0 to *T*.

### Cell culture

Melanoma cell lines were a gift from Dr. Meenhard Herlyn (The Wistar Institute, Philadelphia, PA, USA) and were grown in RPMI-1640 media (Corning, Pittsburgh, PA, USA) supplemented with 5% FBS (Sigma, St. Louis, MO, USA). M257 cells were a gift from Dr. Antoni Ribas (UCLA, Los Angeles, CA, USA).

### Statistical analysis

Matlab statistics toolbox was used to perform the Student’s *t*-test where no significance (ns) denotes > 0.05, ** denotes P < 0.05, and *** denotes P < 0.01.

## Acknowledgments

This study was supported by Moffitt Cancer Center PS-OC NIH/NCI 1U54CA143970-01, R01 CA161107-01, the NCI Comprehensive Cancer Center Grant P30-CA076292, and by monies from the State of Florida to the H Lee Moffitt Cancer Center and Research Institute. The authors thank John Cleveland and Robert Gatenby for critical review of the manuscript.

## Supplementary Materials includes

## Supplementary Materials and Methods

**Figure S1.**
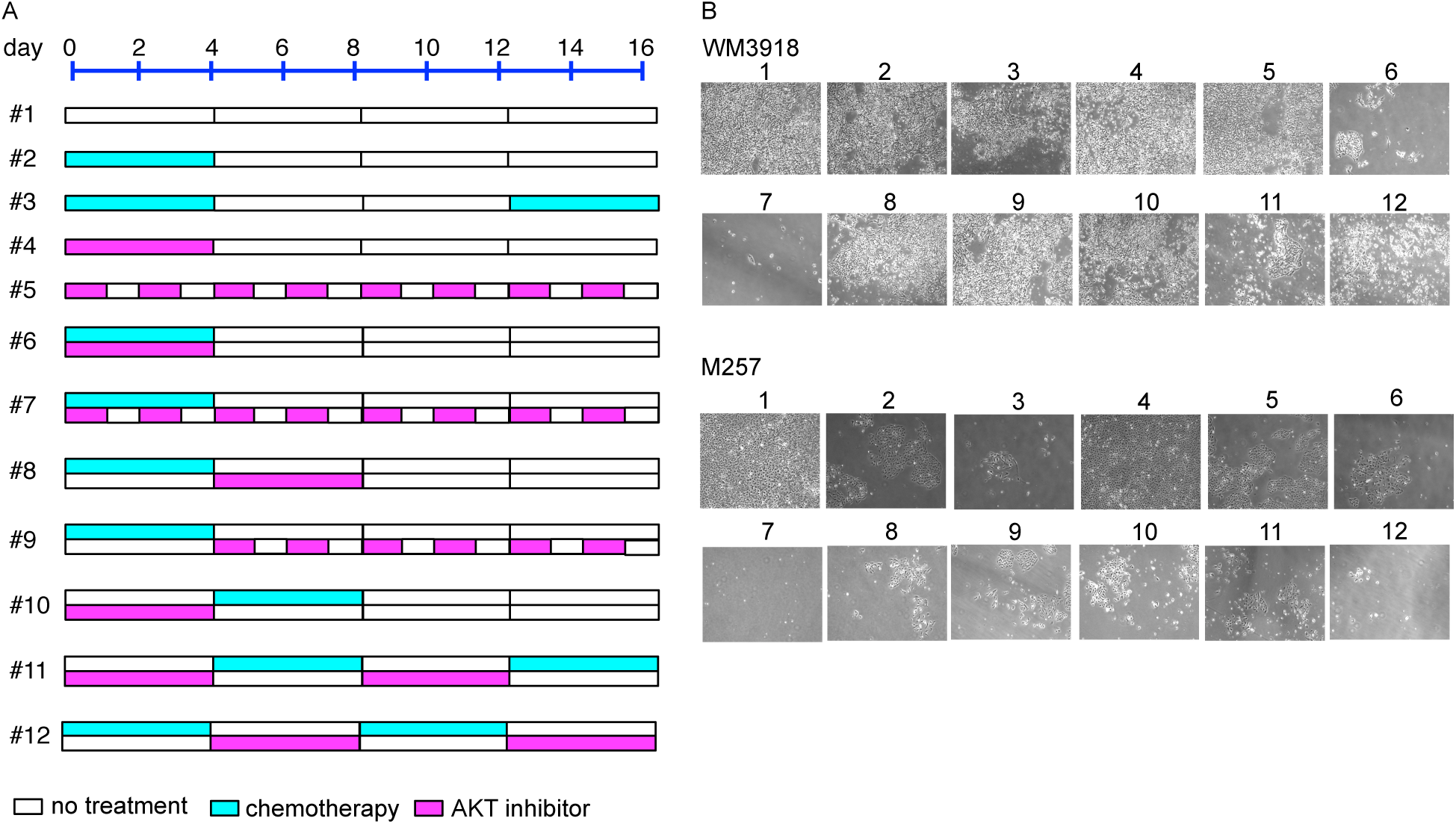
A, graphical representation of the 12 different schedules. B, representative image of cells (WM3918 and M257) on day 16 treated with the schedules. The melanoma cell lines were treated with vehicle, MK-2206 (5 μM), Carboplatin (1 μM), Paclitaxel (3 nM) or the combination of all three agents for 16 days. After this time, colonies were fixed and photographed. Photographs are representative of three independent experiments.

**Figure S2.**
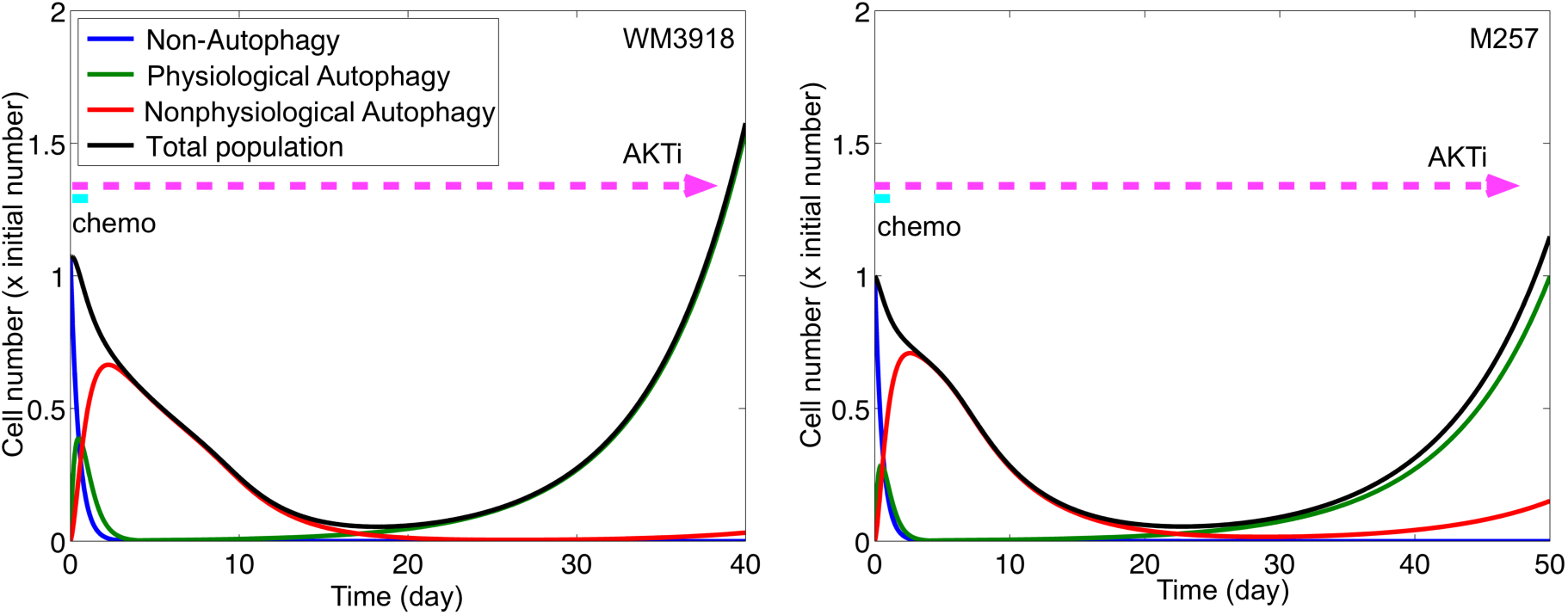
The long-term effects of the therapy (#7).

**Figure S3.**
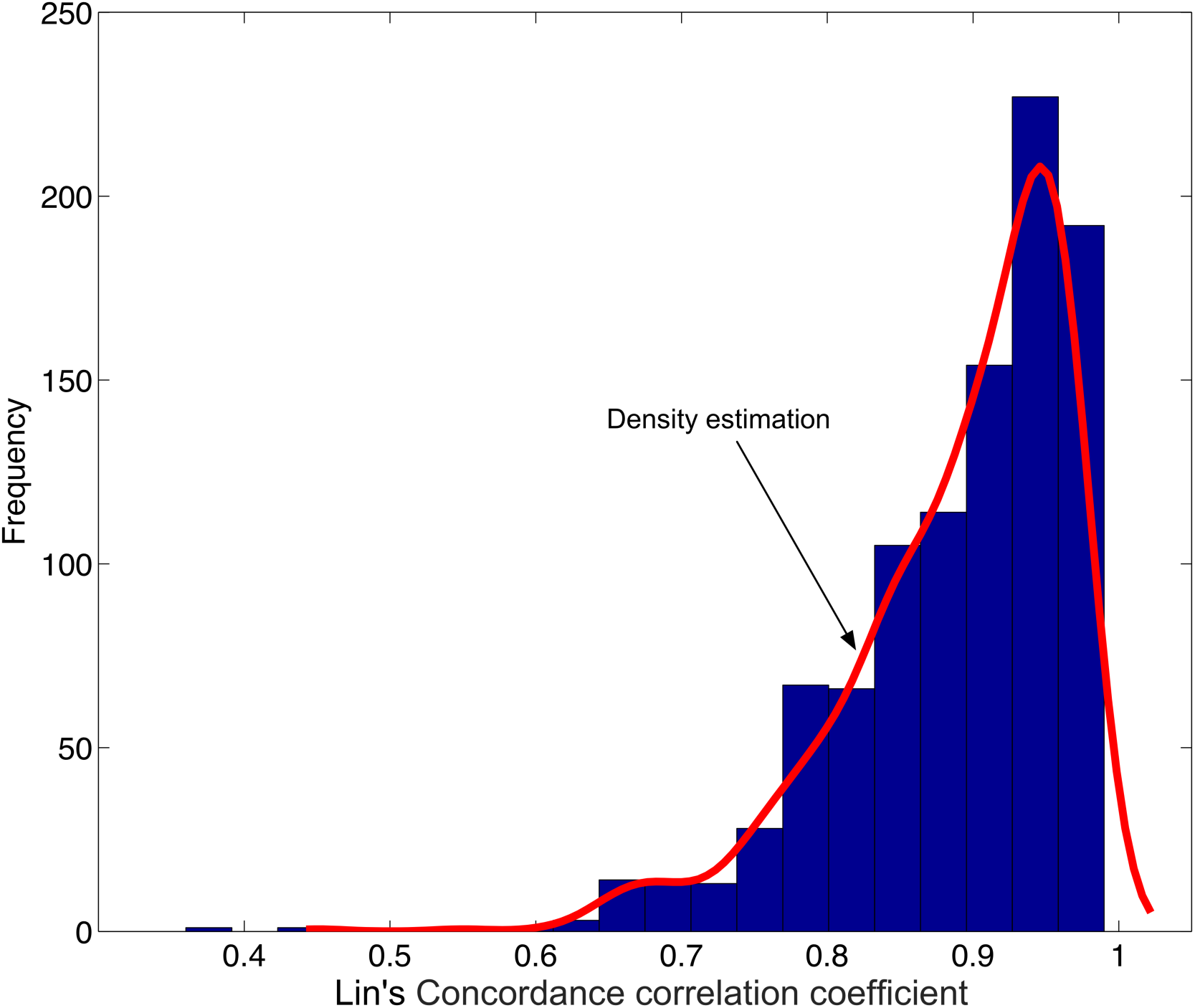
Histogram of Lin’s concordance correlation coefficients and estimated densities based on 1000 samplings from the virtual cohort.

**Figure S4.**
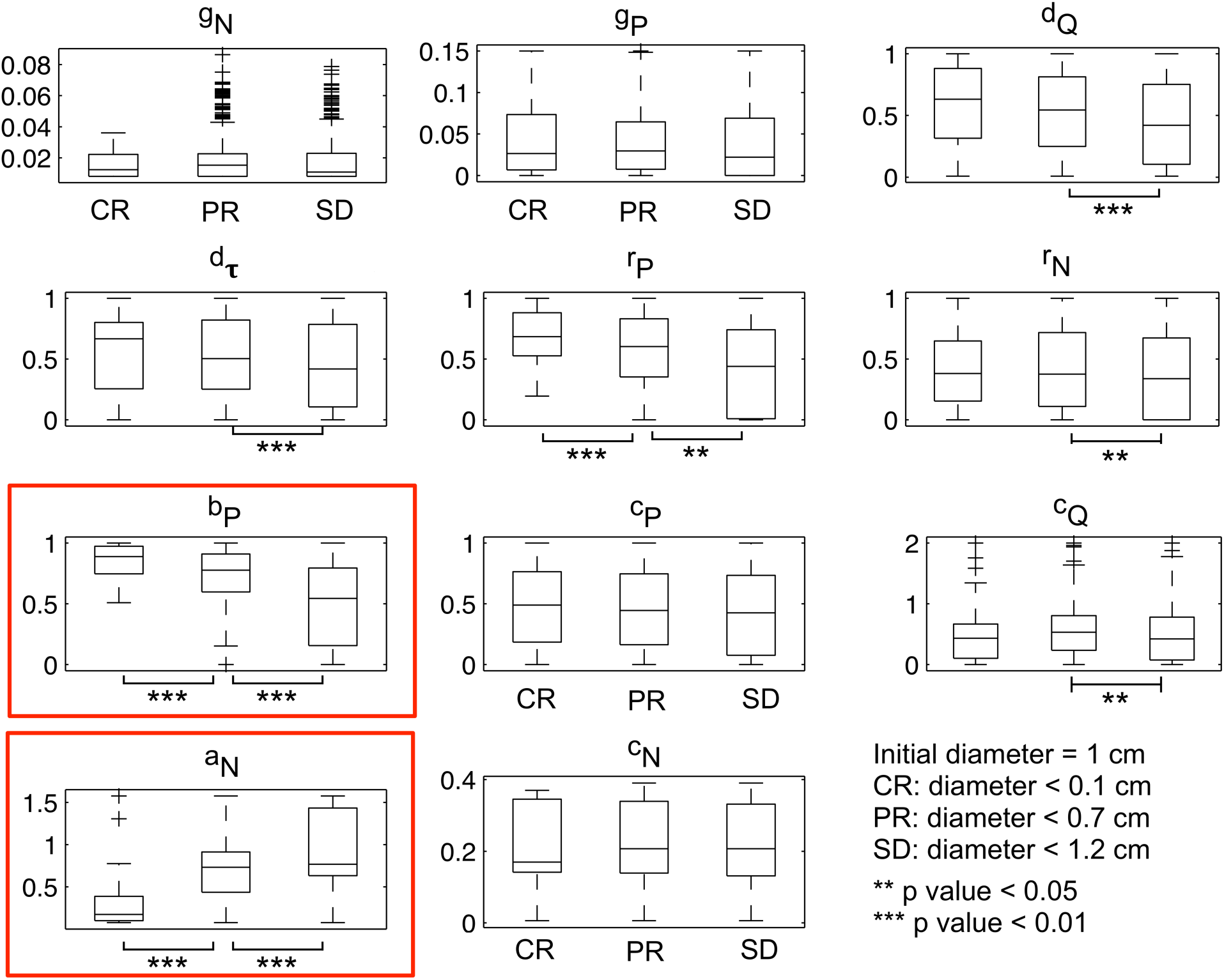
Predictive factors of treatment response arm 1 to the patient cohorts P_1_. For each parameter, a Students’ *t*-test was performed to assess if there is a significant difference in the mean of each model parameter between complete responder (CR), partial responder (PR) and stable diseases (SD). Statistical significance is indicated where ns denotes > 0.05, ** denotes P < 0.05 and *** denotes P < 0.01. Among 11 parameters, *b_P_* and *a_N_* discriminate CR, PR and SD in P_1_ most significantly.

**Figure S5.**
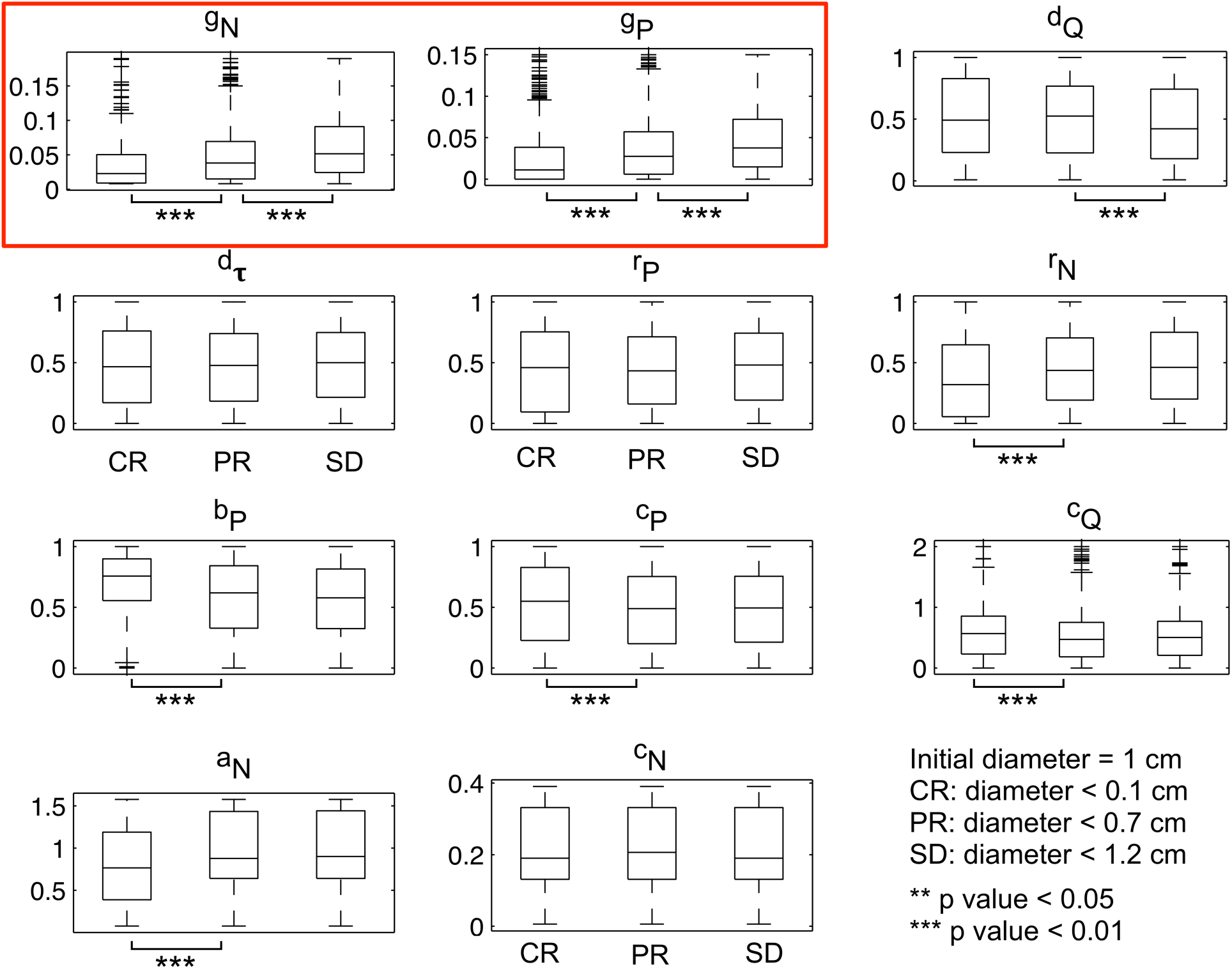
Predictive factors of treatment response arm 2 to the patient cohorts P_2_. Parameter *g_N_* and *g_P_* discriminated CR, PR and SD most significantly.

**Figure S6.**
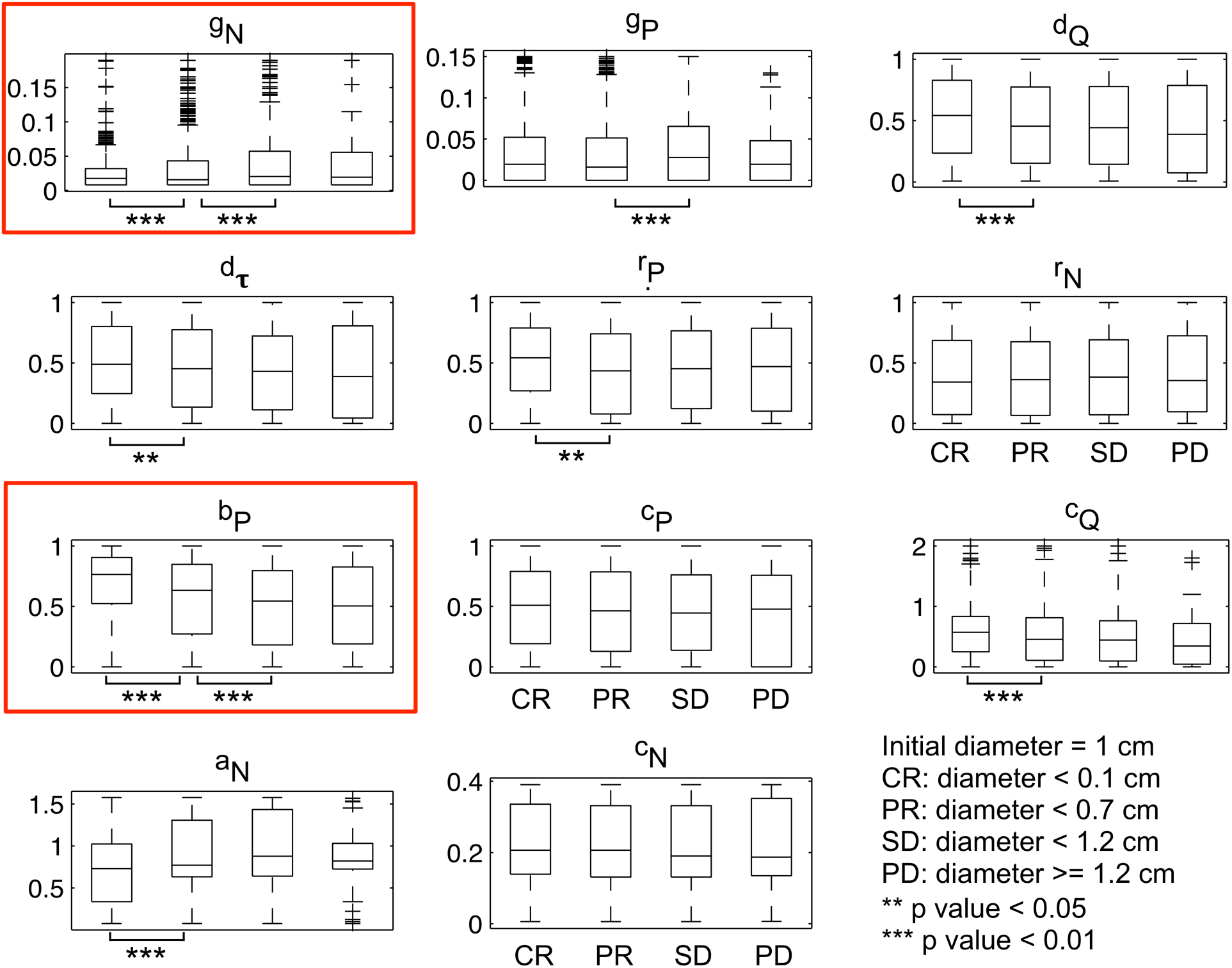
Predictive factors of treatment response either arm 1 or arm 2 to all patients (both P_1_ and P_2_). Parameter *g_N_* and *b_P_* discriminated CR, PR and SD most significantly.

**Table S1.** Estimated parameters (unit: 1/day)

| Name | Description | M257 | WM<br>3918 |
| --- | --- | --- | --- |
| $g_N$ | Growth rate of non-autophagy cells ($N$) | 0.238 | 0.198 |
| $a_0$ | Autophagy transition rate $N \rightarrow P$ | 0.015 | 0.01 |
| $b_0$ | Autophagy transition rate $N \rightarrow Q$ | 0.01 | 0.01 |
| $r_P$ | Recovery rate $P \rightarrow N$ | 0.5 | 0.8 |
| $r_N$ | Recovery rate $Q \rightarrow N$ | 0.7 | 0.5 |
| $g_P$ | Growth rate of physiological autophagy cells ($P$) | 0.19 | 0.195 |
| $q_P$ | Transition rate $P \rightarrow Q$ | 0.05 | 0.01 |
| $r_Q$ | Recovery rate $Q \rightarrow P$ | 0.09 | 0.02 |
| $c_N$ | Increased non-autophagy cell death rate under chemo | 0.3 | 0.1 |
| $c_P$ | Increased physiological autophagy ($P$) cell death rate under chemo | 0.6 | 0.75 |
| $c_Q$ | Increased transition rate $P \rightarrow Q$ under chemo | 1.8 | 1.4 |
| $a_N$ | Increased transition rate $N \rightarrow P$ under AKTi | 1.6 | 2.0 |
| $b_P$ | Increased transition rate $N \rightarrow Q$ under AKTi | 0.15 | 0.15 |
| $d_Q$ | Cell death rate of quiescent autophagy ($Q$) | 0.05 | 0.11 |
| $d_\tau$ | Delayed death rate of quiescent autophagy ($Q$) | 0.15 | 0.2 |
| $\tau$ | Delayed time (unit: day) | 6 | 9 |

A natural death rate ( $d_0$ ) was set to be $d_0 = 0.01$ .

## References and Notes

1. Sawyers C. Targeted cancer therapy. Nature. 2004 Nov 18;432(7015):294–7. PubMed PMID:15549090.

2. Abramson RG. Overview of Targeted Therapies for Cancer. My Cancer Genome2014 [updated September 5, 2014]. Available from: http://www.mycancergenome.org/content/other/molecular-medicine/overview-of-targeted-therapies-for-cancer/.

3. Chapman PB, Hauschild A, Robert C, Haanen JB, Ascierto P, Larkin J, et al. Improved survival with vemurafenib in melanoma with BRAF V600E mutation. The New England journal of medicine. 2011 Jun 30;364(26):2507–16. PubMed PMID: 21639808. Pubmed Central PMCID: 3549296.

4. Arrowsmith J. Trial watch: phase III and submission failures: 2007-2010. Nature reviews Drug discovery. 2011 Feb;10(2):87. PubMed PMID: 21283095.

5. Gan HK, You B, Pond GR, Chen EX. Assumptions of expected benefits in randomized phase III trials evaluating systemic treatments for cancer. Journal of the National Cancer Institute. 2012 Apr 18;104(8):590–8. PubMed PMID: 22491345.

6. Thomas D. Oncology Clinical Trials – Secrets of Success. BIOtechNOW: Biotechnology Industry Organization.; 2012. Available from: http://www.biotech-now.org/business-and-investments/2012/02/oncology-clinical-trials-secrets-of-success.

7. Ledford H. Translational research: 4 ways to fix the clinical trial. Nature. 2011 Sep29;477(7366):526–8. PubMed PMID: 21956311.

8. Lowenstein PR, Castro MG. Uncertainty in the translation of preclinical experiments to clinical trials. Why do most phase III clinical trials fail? Current gene therapy. 2009 Oct;9(5):368–74. PubMed PMID: 19860651. Pubmed Central PMCID: 2864134.

9. Mak IW, Evaniew N, Ghert M. Lost in translation: animal models and clinical trials in cancer treatment. American journal of translational research. 2014;6(2):114–8. PubMed PMID: 24489990. Pubmed Central PMCID: 3902221.

10. Molife LR, Yan L, Vitfell-Rasmussen J, Zernhelt AM, Sullivan DM, Cassier PA, et al. Phase 1trial of the oral AKT inhibitor MK-2206 plus carboplatin/paclitaxel, docetaxel, or erlotinib in patients with advanced solid tumors. Journal of hematology & oncology. 2014;7(1):1. PubMed PMID: 24387695. Pubmed Central PMCID: 3884022.

11. Scott J. Phase i trialist. The Lancet Oncology. 2012 Mar;13(3):236. PubMed PMID:22489289.

12. Clermont G, Bartels J, Kumar R, Constantine G, Vodovotz Y, Chow C. In silico design of clinical trials: a method coming of age. Critical care medicine. 2004 Oct;32(10):2061–70. PubMed PMID: 15483415.

13. Claret L, Girard P, Hoff PM, Van Cutsem E, Zuideveld KP, Jorga K, et al. Model-based prediction of phase III overall survival in colorectal cancer on the basis of phase II tumor dynamics. Journal of clinical oncology: official journal of the American Society of Clinical Oncology. 2009 Sep 1;27(25):4103–8. PubMed PMID: 19636014.

14. Ploquin A, Olmos D, Lacombe D, A’Hern R, Duhamel A, Twelves C, et al. Prediction of early death among patients enrolled in phase I trials: development and validation of a new model based on platelet count and albumin. British journal of cancer. 2012 Sep 25;107(7):1025–30. PubMed PMID: 22910320. Pubmed Central PMCID: 3461164.

15. Holford N, Ma SC, Ploeger BA. Clinical trial simulation: a review. Clinical pharmacology and therapeutics. 2010 Aug;88(2):166–82. PubMed PMID: 20613720.

16. Rebecca VW, Massaro RR, Fedorenko IV, Sondak VK, Anderson AR, Kim E, et al. Inhibition of autophagy enhances the effects of the AKT inhibitor MK-2206 when combined with paclitaxel and carboplatin in BRAF wild-type melanoma. Pigment cell & melanoma research. 2014 May;27(3):465–78. PubMed PMID: 24490764. Pubmed Central PMCID: 3988257.

17. Kondo Y, Kanzawa T, Sawaya R, Kondo S. The role of autophagy in cancer development and response to therapy. Nature reviews Cancer. 2005 Sep;5(9):726–34. PubMed PMID: 16148885.

18. White E. Deconvoluting the context-dependent role for autophagy in cancer. Nature reviews Cancer. 2012 Jun;12(6):401–10. PubMed PMID: 22534666. Pubmed Central PMCID: 3664381.

19. Cheng Y, Zhang Y, Zhang L, Ren X, Huber-Keener KJ, Liu X, et al. MK-2206, a novel allosteric inhibitor of Akt, synergizes with gefitinib against malignant glioma via modulating both autophagy and apoptosis. Molecular cancer therapeutics. 2012 Jan;11(1):154–64. PubMed PMID: 22057914. Pubmed Central PMCID: 3302182.

20. Tormo D, Checinska A, Alonso-Curbelo D, Perez-Guijarro E, Canon E, Riveiro-Falkenbach E, et al. Targeted activation of innate immunity for therapeutic induction of autophagy and apoptosis in melanoma cells. Cancer cell. 2009 Aug 4;16(2):103–14. PubMed PMID: 19647221. Pubmed Central PMCID: 2851205.

21. Liu Y, Levine B. Autosis and autophagic cell death: the dark side of autophagy. Cell death and differentiation. 2015 Mar;22(3):367–76. PubMed PMID: 25257169.

22. Pattingre S, Tassa A, Qu X, Garuti R, Liang XH, Mizushima N, et al. Bcl-2 antiapoptotic proteins inhibit Beclin 1-dependent autophagy. Cell. 2005 Sep 23;122(6):927–39. PubMed PMID: 16179260.

23. Bristol ML, Di X, Beckman MJ, Wilson EN, Henderson SC, Maiti A, et al. Dual functions of autophagy in the response of breast tumor cells to radiation: cytoprotective autophagy with radiation alone and cytotoxic autophagy in radiosensitization by vitamin D 3. Autophagy. 2012 May 1;8(5):739–53. PubMed PMID: 22498493. Pubmed Central PMCID: 3378418.

24. Gewirtz DA. The four faces of autophagy: implications for cancer therapy. Cancer research. 2014 Feb 1;74(3):647–51. PubMed PMID: 24459182.

25. Ma XH, Piao SF, Dey S, McAfee Q, Karakousis G, Villanueva J, et al. Targeting ER stress-induced autophagy overcomes BRAF inhibitor resistance in melanoma. The Journal of clinical investigation. 2014 Mar 3;124(3):1406–17. PubMed PMID: 24569374. Pubmed Central PMCID: 3934165.

26. Goldberg DE. Genetic algorithms in search, optimization, and machine learning. Reading, Mass.: Addison-Wesley Pub. Co.; 1989. xiii, 412 p. p.

27. Holland JH. Adaptation in natural and artificial systems: an introductory analysis with applications to biology, control, and artificial intelligence. Ann Arbor: University of Michigan Press; 1975. viii, 183 p. p.

28. Eisenhauer EA, Therasse P, Bogaerts J, Schwartz LH, Sargent D, Ford R, et al. New response evaluation criteria in solid tumours: Revised RECIST guideline (version 1.1). Eur J Cancer. 2009 Jan;45(2):228–47. PubMed PMID: WOS:000262948300002. English.

29. Burstein HJ, Mangu PB, Somerfield MR, Schrag D, Samson D, Holt L, et al. American Society of Clinical Oncology clinical practice guideline update on the use of chemotherapy sensitivity and resistance assays. Journal of clinical oncology: official journal of the American Society of Clinical Oncology. 2011 Aug 20;29(24):3328–30. PubMed PMID: 21788567.

30. Schrag D, Garewal HS, Burstein HJ, Samson DJ, Von Hoff DD, Somerfield MR, et al. American Society of Clinical Oncology Technology Assessment: chemotherapy sensitivity and resistance assays. Journal of clinical oncology: official journal of the American Society of Clinical Oncology. 2004 Sep 1;22(17):3631–8. PubMed PMID: 15289488.

31. Barth S, Glick D, Macleod KF. Autophagy: assays and artifacts. The Journal of pathology. 2010 Jun;221(2):117–24. PubMed PMID: 20225337. Pubmed Central PMCID: 2989884.

32. Lee HS, Daniels BH, Salas E, Bollen AW, Debnath J, Margeta M. Clinical utility of LC3 and p62 immunohistochemistry in diagnosis of drug-induced autophagic vacuolar myopathies: a case-control study. PloS one. 2012;7(4):e36221. PubMed PMID: 22558391. Pubmed Central PMCID: 3338695.

33. Massey FJ. The Kolmogorov-Smirnov Test for Goodness of Fit. Journal of the American Statistical Association. 1951;46(253):68–78.

34. Bland JM, Altman DG. Statistical methods for assessing agreement between two methods of clinical measurement. Int J Nurs Stud. 2010 Aug;47(8):931–6. PubMed PMID: WOS:000280343000002. English.

35. Lin LI. A concordance correlation coefficient to evaluate reproducibility. Biometrics. 1989 Mar;45(1):255–68. PubMed PMID: 2720055.

36. Stevenson M. Tools for the Analysis of Epidemiological Data. 0.9-62 ed2015.

37. Carlson JA. Tumor doubling time of cutaneous melanoma and its metastasis. The American Journal of dermatopathology. 2003 Aug;25(4):291–9. PubMed PMID: 12876486.

38. Kelley CT. Implicit filtering. Philadelphia: Society for Industrial and Applied Mathematics; 2011. xiv,170 p. p.

39. Das Thakur M, Salangsang F, Landman AS, Sellers WR, Pryer NK, Levesque MP, et al. Modelling vemurafenib resistance in melanoma reveals a strategy to forestall drug resistance. Nature. 2013 Feb 14;494(7436):251–5. PubMed PMID: 23302800. Pubmed Central PMCID: 3930354.

40. Leder K, Pitter K, Laplant Q, Hambardzumyan D, Ross BD, Chan TA, et al. Mathematical modeling of PDGF-driven glioblastoma reveals optimized radiation dosing schedules. Cell. 2014 Jan 30;156(3):603–16. PubMed PMID: 24485463. Pubmed Central PMCID: 3923371.

41. Ohtsu T, Sasaki Y, Tamura T, Miyata Y, Nakanomyo H, Nishiwaki Y, et al. Clinical pharmacokinetics and pharmacodynamics of paclitaxel: a 3-hour infusion versus a 24-hour infusion. Clinical cancer research: an official journal of the American Association for Cancer Research. 1995 Jun;1(6):599–606. PubMed PMID: 9816021.

42. Siddiqui N, Boddy AV, Thomas HD, Bailey NP, Robson L, Lind MJ, et al. A clinical and pharmacokinetic study of the combination of carboplatin and paclitaxel for epithelial ovarian cancer. British journal of cancer. 1997;75(2):287–94. PubMed PMID: 9010040. Pubmed Central PMCID: 2063279.

43. Yap TA, Yan L, Patnaik A, Fearen I, Olmos D, Papadopoulos K, et al. First-in-man clinical trial of the oral pan-AKT inhibitor MK-2206 in patients with advanced solid tumors. Journal of clinical oncology: official journal of the American Society of Clinical Oncology. 2011 Dec 10;29(35):4688–95. PubMed PMID: 22025163.

